# Identifiability-Guided Assessment of Digital Twins in Alzheimer’s Disease Clinical Research and Care

**DOI:** 10.1101/2025.08.17.670697

**Authors:** Juliet Jiang, Jeffrey R. Petrella, Wenrui Hao, the Alzheimer’s Disease Neuroimaging Initiative

## Abstract

Digital twins – personalized, data-driven computational models – are emerging as a powerful paradigm for representing and predicting disease trajectories at the individual level. These models have the potential to support diagnosis, monitor disease evolution, and evaluate therapeutic interventions in virtual settings in the context of clinical trials and patient care. Rigorous model assessment is thus critical for its implementation, but medical data are often sparse, noisy, and vary significantly across individuals, making it challenging to determine whether a digital twin optimized on such data is valid. In such settings, identifiability analysis becomes essential for evaluating whether model parameters can be reliably estimated and interpreted. To address this, we investigate how identifiability can support the clinical application of a computational causal digital twin model for Alzheimer’s Disease (AD), where data sparsity and variability are particularly pronounced. Our results show that the magnitude and distribution of biomarker data influence the parameter practical identifiability, and that constraints on the model structure and parameters can significantly affect identifiability. We also observe differences in identifiability across diagnostic groups, with several parameters showing significantly different values between individuals with AD, mild cognitive impairment (MCI), and cognitively normal (CN) subjects. Uncertainty quantification for identifiable parameters and their corresponding model trajectories provides visual insight into variability in disease progression and reveals mild trends related to biomarker data spread. This study represents a first step toward incorporating identifiability techniques into clinical digital twin frameworks, using a data-driven, interpretable example based on a previously published AD model.

## Introduction

Alzheimer’s disease (AD) is a chronic, progressive neurodegenerative disorder and the most common cause of dementia in older adults. It is characterized by a prolonged preclinical phase, heterogeneous symptom onset, and complex interactions among biological pathways—including amyloid accumulation, tau pathology, neurodegeneration, and cognitive decline. Despite decades of research, effective therapeutic strategies remain limited, partly due to the disease’s long timescale, significant inter-individual variability, and diagnostic uncertainty. Clinical categories such as mild cognitive impairment (MCI) and Alzheimer’s dementia provide only coarse snapshots of disease status and often fail to capture the dynamic, individualized nature of disease progression^1–3^. Disease subtyping, such as the Amyloid Tau Neurodegeneration (ATN) framework in AD, based on binary biomarker categories, is a first step toward accounting for inter-individual variability and reducing diagnostic uncertainty. However, there still remains considerable variability in the clinical course and therapeutic response within ATN categories^4–6^.

As precision medicine becomes a central goal in AD care, there is an urgent need for tools that integrate patient-specific biomarker data to guide individualized prediction and treatment. Computational modeling offers a promising path toward this goal by providing mechanistic insights into disease progression and enabling simulation-based forecasting. In particular, digital twins—personalized, data-driven computational models—are emerging as a powerful paradigm for representing and predicting disease trajectories at the individual level^7,8^. These models have the potential to support diagnosis, monitor disease evolution, and evaluate therapeutic interventions in virtual settings.

Over the past two decades, a wide range of mathematical and computational models have been developed to characterize AD biomarker progression, varying in mechanistic complexity and degree of personalization. Early models focused on the aggregation kinetics of amyloid-*β* and tau proteins, employing systems of differential equations to describe their production, clearance, and interactions. Subsequent models incorporated neurodegeneration and cognitive decline, often embedding nonlinear feedback mechanisms among biomarkers to better reflect disease complexity^9,10^. Spatially explicit, network-based models have been used to simulate the propagation of pathology through the brain’s structural connectome^11–13^. Parallel to these mechanistic approaches, statistical and machine learning models have enabled data-driven characterizations of disease progression. Data-driven algorithms can predict trajectories with high accuracy, but the application of computational models to clinical settings requires that models remain mechanistically grounded in order to have accurate uncertainty quantification and physical interpretability^8^. Latent time models, event-based frameworks, and disease progression scoring methods have been applied to estimate disease staging and predict future biomarker trajectories^14^. These studies range from simulating the disease trajectory of a single subject^9,12^ or mapping a general disease progression through regression^14^, to optimizing individual treatment therapies in dozens of patients^10,13^ or modeling interactive biomarker mechanisms in hundreds^11,15^.

A computational causal model of AD based on the Alzheimer’s Disease Biomarker Cascade (ADBC) theory—a widely recognized framework for AD progression—has been validated against clinical data as a foundational step toward developing AD digital twins^15^. The fitting of such models depends on accessible and reliable clinical data, yet traditional clinical trials face numerous limitations, including high costs, long durations, limited sample sizes, data entry errors, and ethical concerns^16^. Moreover, clinical datasets are frequently incomplete or collected under variable experimental conditions, complicating reliable parameter estimation^17–19^. Although computational models and *in silico* simulations leveraging digital twins offer promising solutions to these challenges^20–22^, they often involve many parameters relative to typically sparse and noisy patient data. As computational models gain traction in clinical and regulatory contexts—such as digital clinical trials—the issue of ambiguous parameter estimation becomes critical for model validity and practical utility^23^. Poorly determined parameters lead to uncertain predictions, undermining confidence in model-driven decisions. Conversely, when parameters cannot be reliably inferred from available data, it is essential to reduce costs and patient burden by avoiding noninformative measurements^24^.

*Identifiability* analysis has emerged as a crucial tool for assessing the feasibility and reliability of parameter estimation, especially in complex or high-dimensional dynamical systems^24–26^. Previous work highlights that identifiability can be used to reduce the cost of data collection and provide insight into how we can enable better incorporation of mathematical models into clinical applications^12,24^. Parameters are deemed *non-identifiable* when available experimental data are insufficient relative to model complexity. Even identifiable parameters yield estimates accompanied by uncertainty, typically expressed as *confidence intervals* that quantify the probable range of true parameter values^27^. Structural identifiability examines whether a unique parameter set theoretically exists based on model structure alone, independent of data quality, whereas practical identifiability considers the influence of data quantity and quality on parameter estimation^27–29^.

In this work, we investigate the practical identifiability of nine parameters in the mechanistic ADBC model introduced previously by our group^15^. Our goals are to provide clinicians with interpretable and informative insights specific to our AD model and to emphasize the importance of identifiability analysis in future disease modeling efforts aimed at clinical translation. We assess practical identifiability across a population of subjects, analyzing its relationship to data characteristics and disease states, and utilize these results to inform a downstream uncertainty quantification. A deeper understanding of identifiability and a more reliable presentation of predictive uncertainty in clinical contexts can promote its integration into experimental design, strengthen its role in clinical decision-making, and highlight the complexities inherent in biologically interpretable models beyond theoretical constructs.

## Results

### Parameter identifiability is associated with spread or magnitude

Parameter identifiability is determined by whether the profile likelihood-based confidence interval of a parameter is finite. Assessments of the profile likelihoods (Figure 2a) for each subject provide unique identifiability profiles across the cohort. Unique parameter estimation is naturally impossible if the number of data points fails to reach the degrees of freedom required to constrain the model, that is, when the number of observations is less than the number of independent parameters to be estimated. Including the constraint of reaching a maximal disease state at age 100, sigmoidal curves require a minimum of two additional points. All subject data have at least two data points in amyloid-*β*, phosphorylated tau, and neurodegeneration. However, clustered data points can operate virtually as one data point, suggesting that the number of data points alone is not a sufficient indicator of practical identifiability. We randomly generated 100 synthetic data points from several disease trajectories and re-evaluated parameter identifiability to check this, and found that even with an overabundant number of data points, there was not a single case where all nine parameters were identifiable. Therefore, we seek to assess the quality of modeling data as well. We observe quality through the vertical and horizontal spread of data and data magnitude, captured through metrics such as the mean and standard deviation of both biomarker data and ages of examination. From the logistic regressions in Figure 2c, we observe that all six metrics (mean, median, standard deviation, range, maximum, and minimum) influenced the identifiability of 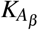 (carrying capacity of amyloid), and the mean, median, and max/min largely influenced the identifiability of *A*_0_ (initial amount of amyloid). Decision boundaries were less clear for 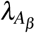 (growth rate of amyloid) and for parameters associated with tau and neurodegeneration, which are listed in the supplementary. Logistic regressions against similar metrics for ages during examinations are also in the supplementary and show no clear trends.

**Figure 1.**
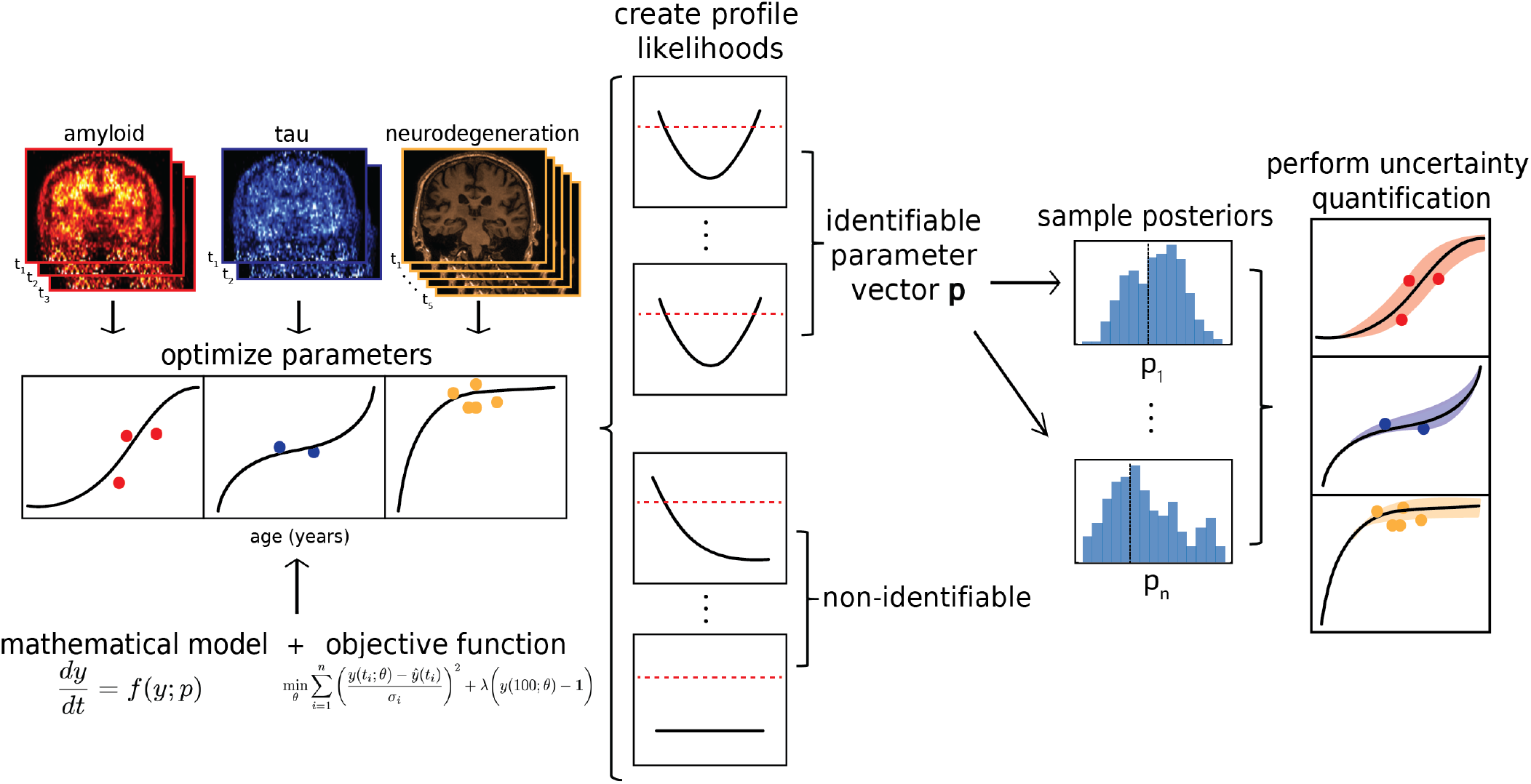
Schematic diagram of the proposed computational modeling framework. PET scan SUVR measurements from summary regions of interest (ROIs) in a cohort of 223 subjects represented amyloid-*β* and phosphorylated tau biomarkers. Hippocampal volume, normalized by intracranial volume (ICV), represented the neurodegeneration biomarker. A system of differential equations was fitted to individual patient data via an objective function. Profile likelihoods were then computed around optimal parameter estimates to assess practical identifiability. Joint posterior distributions of identifiable parameters were sampled using Monte-Carlo Markov chain sampling, enabling the construction of a 95% confidence interval around the average disease trajectory.

**Figure 2.**
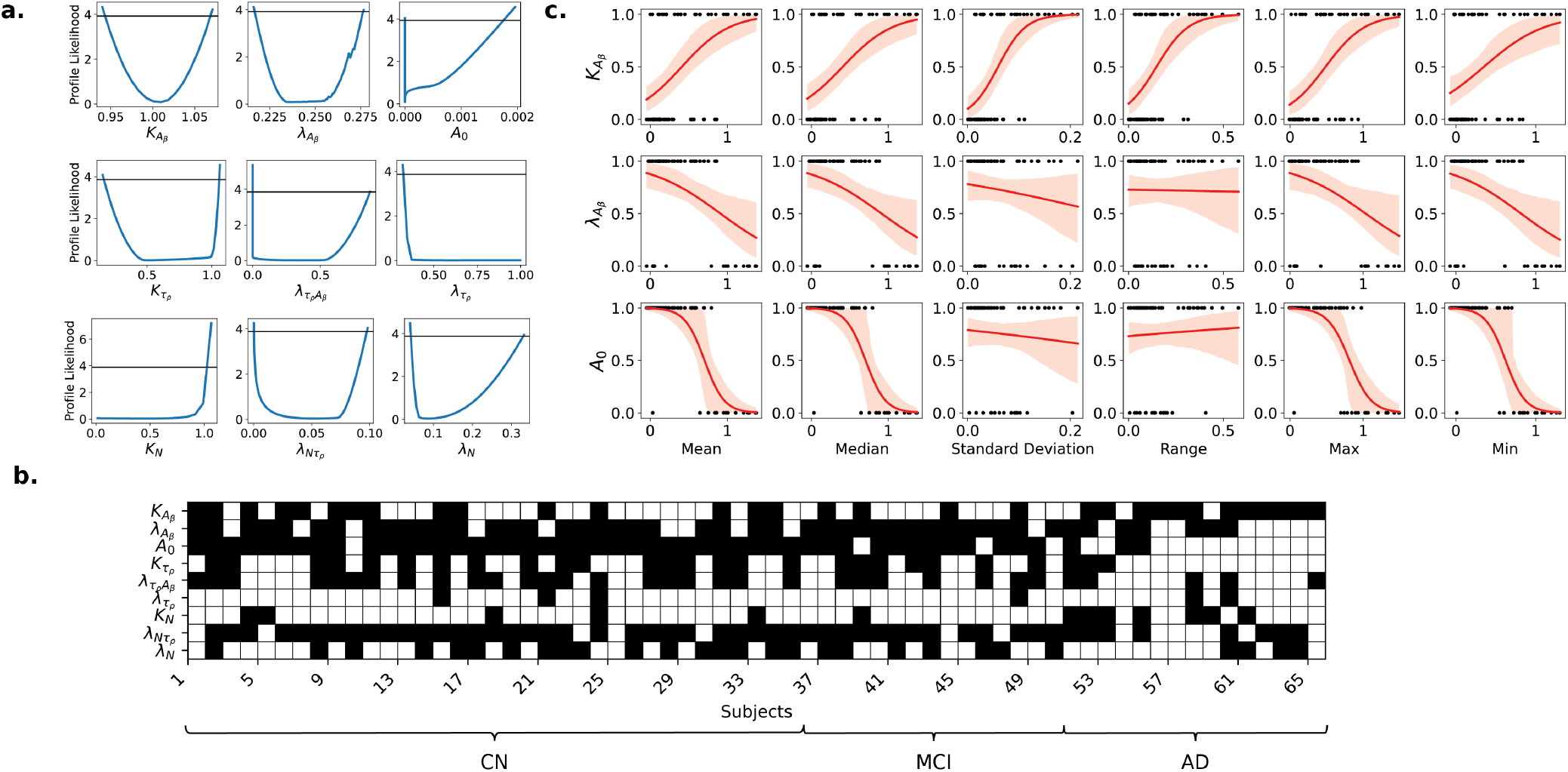
a. Profile likelihoods of all nine parameters for an example subject. b. Binary map of practical identifiability for all nine parameters across 65 (35 cognitively normal (CN), 15 MCI, 15 AD) of the 223 subjects, grouped by clinical diagnosis. Black - identifiable, white - non-identifiable. c. Logistic regressions for the three amyloid-related parameters (corresponding tau and neurodegeneration plots are in the supplementary material) against the mean, median, standard deviation, range, maximum, and minimum of amyloid data for each subject.

### Identifiability varies across different diagnostic groups

While the spread and magnitude of data govern a subject’s endophenotypic profile, these profiles may not match clinically diagnosed disease labels^15^. We are also interested in how identifiability may evolve across disease states – these results are summarized visually in Figure 2c and numerically in Table 1. The identifiability of 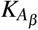 and *A*_0_ are significantly different across diagnostic groups, with both omnibus significances driven by large differences in the AD group (Table 2). Identifiability of 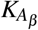 is higher in AD subjects, while a decrease in the identifiability of *A*_0_ can be observed from the CN to AD state. We also found that 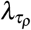 and *K*_*N*_ are the least identifiable parameters among the nine (IF of 0.092 and 0.200, respectively), but both parameters show the largest percentage of identifiability in the AD group. Other rate parameters such as 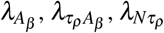, and *λ*_*N*_ see a decline in identifiability from CN to AD. Parameters for amyloid-*β* are, at large, more identifiable than parameters for phosphorylated tau.

**Table 1.** Identifiability fractions (IF) across diagnostic groups.

| Parameter | IF (CN) | IF (MCI) | IF (AD) | Overall IF |
| --- | --- | --- | --- | --- |
| $K_{A_\beta}$ | 0.4286 | 0.2667 | 0.8000 | 0.477 |
| $\lambda_{A_\beta}$ | 0.7429 | 0.9333 | 0.4667 | 0.723 |
| $A_0$ | 0.9714 | 0.8000 | 0.2000 | 0.754 |
| $K_{\tau_p}$ | 0.4857 | 0.4667 | 0.2000 | 0.415 |
| $\lambda_{\tau_p A_\beta}$ | 0.6000 | 0.4667 | 0.3333 | 0.508 |
| $\lambda_{\tau_p}$ | 0.0857 | 0.0667 | 0.1333 | 0.092 |
| $K_N$ | 0.1429 | 0.0667 | 0.4667 | 0.200 |
| $\lambda_{N\tau_p}$ | 0.8571 | 0.8667 | 0.5333 | 0.785 |
| $\lambda_N$ | 0.4857 | 0.4667 | 0.2667 | 0.431 |

**Table 2.** Significance results from an omnibus ANOVA test for each parameter and pairwise post-hoc tests.

|  | Omnibus | CN vs MCI | CN vs AD | MCI vs AD |
| --- | --- | --- | --- | --- |
| $K_{A_\beta}$ | 0.004585* | 0.914991 | 0.019897* | 0.006936* |
| $\lambda_{A_\beta}$ | 0.019727 | 0.227272 | 0.428213 | 0.014850* |
| $A_0$ | 0.000004* | 0.588237 | 0.000008* | 0.000182* |
| $K_{\tau_p}$ | 0.149858 | 0.227272 | 0.428213 | 0.014850 |
| $\lambda_{\tau_p A_\beta}$ | 0.194090 | 0.518976 | 0.170876 | 0.744942 |
| $\lambda_{\tau_p}$ | 0.358942 | 0.750402 | 0.325531 | 0.750402 |
| $K_N$ | 0.033164 | 0.649410 | 0.188501 | 0.028329 |
| $\lambda_{N\tau_p}$ | 0.049107 | 1.000000 | 0.082004 | 0.082004 |
| $\lambda_N$ | 0.321290 | 0.928612 | 0.316578 | 0.518976 |

### Uncertainty quantification using posteriors of identifiable parameters

Uncertainty quantification allows us to assess the limitations of a model’s predictive power, but should only be performed with identifiable parameters^30–32^. After estimating the posterior distributions with MCMC sampling, we drew 1,000 samples to simulate a range of plausible disease progression trajectories, each corresponding to a different parameter set. By averaging these simulated trajectories and computing the 95% credible intervals, we visualized the uncertainty in an individual’s predicted disease profile, as illustrated in Figure 3a. The confidence intervals at each data point were averaged for each subject and plotted against the same quantitative metrics, shown in Figure 3b. Trends indicate that data points with the least standard deviation and magnitude have the tightest confidence intervals. Linear regression on each scatterplot in Figure 3b reveals that the highest slopes were found between the averaged confidence intervals for amyloid and spread metrics like standard deviation and range (Figure 3c). These two relationships were also the only two metrics that showed significance in the hypothesis tests, with p-values at 0.01451 and 0.01758, respectively.

**Figure 3.**
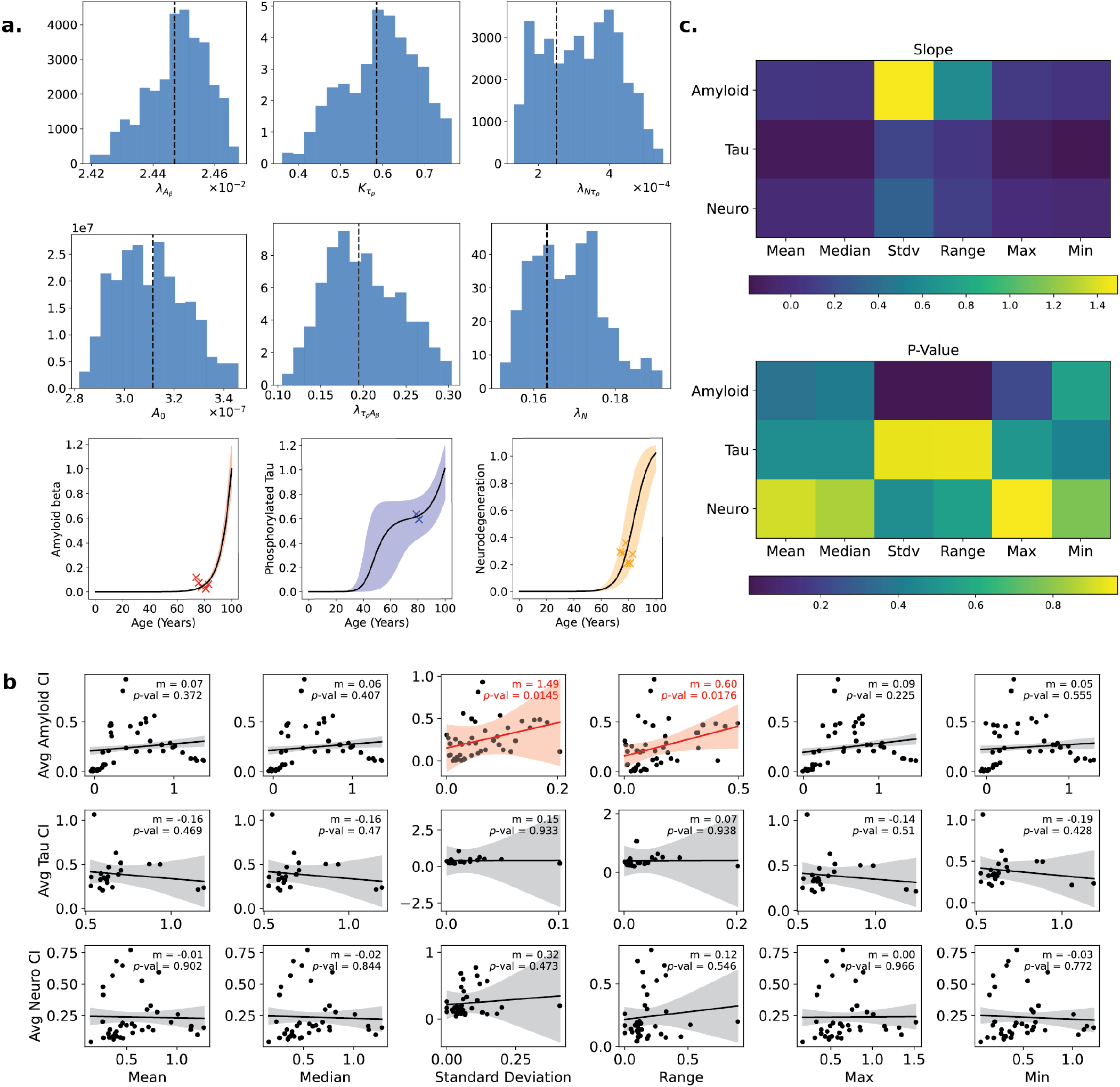
a. Uncertainty quantification from perturbing identifiable parameters within the 95% confidence bounds of their joint posterior distribution in an example subject. In columns from left to right: amyloid-*β*, phosphorylated tau, and neurodegeneration. b. Averaged confidence interval widths of predicted disease progression across data points are plotted against the mean, median, standard deviation, range, maximum, and minimum of biomarker data (n = 45). c. Slope and p-values obtained from a linear regression performed on each scatterplot in panel b.

## Discussion

Digital twins have the potential to play a transformative role in personalized medicine by enabling patient-specific predictions and treatment planning. Trust in these digital representations depends on accurate and reliable mathematical models. In real-world settings where medical data are sparse, noisy, and heterogeneous, it becomes essential to evaluate how well model parameters can be realistically estimated and whether the resulting simulations can be interpreted with confidence. In this study, we used longitudinal data in individual subjects drawn from a widely recognized database in Alzheimer’s research. We used identifiability analysis as a guide to determine the reliability of digital twin simulations of individual disease progression scenarios, and to assess what characteristics of the data predict high identifiability, and therefore clinically reliable simulations.

Regarding characteristics of the data that predict high identifiability, we found that the number of data points available is not significantly related to identifiability, with data magnitude and spread serving as more important features in our model and against our model assumptions. Parameters such as 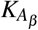, the carrying capacity for amyloid, are more likely to be identifiable with a larger standard deviation and magnitude in amyloid data, which corresponds to a more defined slope and data points closer to the constraint at age 100 (Equation 4). Low magnitude amyloid data tended to cluster more than high magnitude amyloid data, as those with amyloid accumulation that was originally high continued to increase, while those without accumulation remained healthy. Trajectories fitted to amyloid data clustered near zero lack an inflection point and show no evidence of approaching a carrying capacity within the observed time frame, leading to frequent non-identifiability. Conversely, *A*_0_, the initial value of amyloid, was more likely to be identifiable with lower amyloid values. All parameters were optimized using a bound constrained algorithm, with biologically plausible parameter bounds set to those in Table 3. The assumption that amyloid-*β* aggregation is minimal at birth means that a subject’s optimal amyloid trajectory may initialize very close to this upper bound. Without an enclosed likelihood-based confidence interval, the parameter becomes non-identifiable within the biologically realistic range. Trade-offs between model assumptions and non-identifiability due to these assumptions and constraints have been noted in other papers^33,34^, but should be further explored and generalized. Additionally, the vertical spread of the biomarkers accounts more for the variability in slope because the horizontal spread in subject ages is relatively fixed from evenly scheduled examination dates. Variation in ages of examination is also typically low, less than 10% of the entire modeled lifespan. Analysis done on data collected from a larger span of years with longer longitudinal follow up could provide greater insight into the impact of data collection frequency and consistency.

**Table 3.** Optimization bounds for each model parameter used during parameter estimation.

| Parameter | Lower Bound | Upper Bound |
| --- | --- | --- |
| $K_{A\beta}$ | 0 | 10 |
| $\lambda_{A\beta}$ | 0 | 1 |
| $A_0$ | 0 | 0.2 |
| $K_{\tau_p}$ | 0 | 10 |
| $\lambda_{\tau_p A\beta}$ | 0 | 1 |
| $\lambda_{\tau_p}$ | 0 | 1 |
| $K_N$ | 0 | 10 |
| $\lambda_{N\tau_p}$ | 0 | 1 |
| $\lambda_N$ | 0 | 1 |

These patterns are reflected in differences in parameter identifiability across diagnostic groups. Many subjects with AD possessed levels of amyloid-*β* data, and in some cases neurodegeneration data, above the constraint at (100,1), leading to carrying capacities 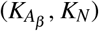 that were significantly more identifiable than those in CN and MCI subjects. Fitting data above the constraint necessitates that trajectories meet the carrying capacity before age 100, and these carrying capacity parameters will have to be optimized to near 1. These same patterns in high amyloid aggregation and greater neurodegeneration led *A*_0_ and the growth constants for amyloid and neurodegeneration 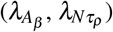 to be significantly less identifiable in AD subjects. This is because biomarker data were collected later in the given lifespan, so those with high magnitude biomarker values have sigmoid-shaped trajectories with arbitrarily high slope before the first examination. If the rate constants can be arbitrarily large due to a lack of slope-determining data, they become non-identifiable. And again, non-identifiability of *A*_0_ in AD subjects can also be attributed to the likelihood-based confidences intervals exceeding the range of biologically plausible parameter values. Subjects in the CN group often had low amyloid aggregation, leading to significantly more identifiable *A*_0_ parameters.

To our knowledge, no practical identifiability study exists for AD models. Rahimabadi and Benali^35^ assess the structural identifiability of a model for tauopathy progression in the mouse brain using reaction–diffusion equations, but practical identifiability is missing. Nonetheless, findings from prior cancer and infectious disease models remain applicable and can meaningfully inform our work. In the realm of complex, progressive diseases with multifactorial origins, the literature is rich with practical identifiability studies on cancer models^34,36–39^ and infectious disease models^40–43^. Some of the cancer studies also use clinical data^33,44^. Phan et al.^44^ also studied model identifiability from a clinical perspective, noting the frequency of data collection, the types of data, and the accuracy of measurements as contributors to the practical identifiability of their model. Using synthetic data and an observing system simulation experiment (OSSE) framework to determine identifiability, they found that higher frequency of data collections greatly increased identifiability, and that data should cover multiple temporal regions of cancer growth. Saucedo et al. presented similar conclusions on the importance of data frequency and data type in fitting an SEIR model, using Monte Carlo simulations and a Correlation Matrix approach with synthetic data^43^. Similarly, our work found features beyond just the number of data points to be critical, but extended beyond temporal characteristics while acknowledging that identifiability would benefit from a greater temporal range of data. Eisenberg and Jain^37^ evaluated non-identifiability in two chemotherapy models, finding biological insights in practically identifiable parameter combinations. Their work (and others^26,45^) highlights the link between non-identifiability and optimal experimental design, and found different data collection strategies that reduced cost while maintaining the same model uncertainty. In contrast, our work is primarily a post hoc analysis using existing ADNI data, aimed at exploring how identifiability and uncertainty can be assessed a priori based on the available data. With greater temporal variability in our data, our approach could offer additional guidance for optimizing future clinical examination schedules. These authors also discuss the necessary collaboration between modelers and experimentalists to generate tailor-made models and data. These goals may still be idealistic in AD models since cancers progress much faster than Alzheimer’s, making this iterative modeling process drastically slower.

Another strength of this work is the extension of identifiability analysis on individual subjects to the population level. Previous work presents case studies that are limited by small datasets. With personalized models and treatment being the target application of our model, separate identifiability analyses for each subject present the idea of *personalized identifiability profiles* alongside individualized models. The ability to have identifiability profiles across a population of subjects enables us to also analyze differences in identifiability across disease subgroups, and determine group-specific variability in parameter estimability. Our work is the first to explicitly use practical identifiability on patient-specific medical data and extract disease-related outcomes.

Other relationships with data characteristics emerged from our study of uncertainty. For example, the confidence intervals of the model solutions for amyloid-*β* had a positive linear relationship with the vertical spread of biomarker data. This relationship was less clear for the horizontal spread of datapoints, likely due to little variability in age. In general, confidence intervals highly varied across subjects and across biomarkers (e.g. Figure 3a). Some were consistently very narrow and some were wider through the later half of a subject’s lifespan, while others depicted low confidence prior to age 70 and tightened in the presence of data, a phenomenon most frequently observed in the tau trajectories. Subjects with low amyloid-*β* values, often the case in CN subjects, tended to have very small confidence intervals. This has clinical implications, as intervention to change disease progression is best administered, as early as possible, between the ages 60-85^46,47^. Model predictions may provide high confidence, in the asymptomatic or prodromal stages of disease, that biomarker progression will drastically accelerate or remain slow in the next decade of life. These types of projections, coupled with genetic risk factors such as the Apolipoprotein E (APOE) *ε*4 allele^48,49^, underscore the importance of uncertainty quantification for accurate risk prediction and clinical decision-making, guiding both the urgency and choice of intervention as well as the development of appropriate therapies^8^.

For clinicians to use digital twins in decision-making, it is essential for digital twins to have carefully communicated prediction uncertainty under rigorous methods and have it be tightly integrated in the modeling process. Uncertainty quantification will play a deeper role in the design and deployment of digital twins, requiring standardized methods that can adapt to data, rather than just model structure^8,50,51^. Being the first practical identifiability study on AD models, our work is also the first to employ identifiability to assess predictive model uncertainty in AD models. Corti et al.^12^ constructed an uncertainty quantification framework and applied it to a patient-specific model for amyloid-*β* accumulation across brain regions. They used MCMC sampling without prior identifiability analysis to obtain posterior distributions, but MCMC sampling in the presence of non-identifiability can be misleading^32^. It is important that uncertainty quantification is applied only to identifiable parameters, as uncertainties propagating from an unidentifiable model can be weak in reproducibility and stability^30,31^. Previous uncertainty quantification on AD models relied on model sensitivity, without the consideration of data quality and quantity. An identifiability-informed, data-driven uncertainty quantification framework at the individual level is necessary for the development of and trust in digital twins.

This work has a number of limitations. The maximum number of medical data points per biomarker is capped at seven, with most measurements concentrated within a single decade of life, making reliable parameter estimation inherently challenging. Of the 800+ subjects in the broader dataset, only 223 had at least two data points for all three biomarkers, and all of these had just two or three tau scans. Furthermore, because the earliest examination begins at age 55, the first half of each subject’s lifespan is entirely data-deficient. Although the constraint anchoring biomarkers at time = 0 (excluding amyloid) and time = 100 years facilitated initial optimization, it occasionally introduced numerical instability during profile likelihood computations. Future models can be restricted to the last few decades of the lifespan and utilize a loosened constraint. Likewise, difficulties in sampling convergence, numerically stable profile likelihoods, and finding appropriate model solutions in the initial optimization required copious hyperparameter tuning, such as the step size and the number of alternations between parameters in coordinate MCMC sampling, the constraint coefficient in the objective function, the scanning resolution and bounds for finding the profile likelihood, and more. Excessive hyperparameter tuning makes model generalization weak, and future work can involve the use of other methods, such as adaptive step sizes and no constraint, to reduce the number of hyperparameters. Moreover, dependencies between biomarkers within the cascading nature of the ADBC model are unclear. For example, few tau parameters were identifiable, so practical identifiability across biomarkers was not mathematically formalized or explained. Few tau parameters were identifiable potentially for these unknown dependencies, but much more likely is the sparsity of tau data. We lack understanding in why neurodegeneration-related parameter identifiability is still low despite having a high number of data points, but as neurodegeneration follows tau in the cascade hypothesis, it is possible non-identifiability of the former is leading to flaws in the estimation of the downstream variable. Little work exists in investigating how practical identifiability evolves over sequentially solved equations, as most model parameters are optimized simultaneously. Therefore, while 2D profile likelihoods^52,53^ and parameter paths^54^ can normally be very helpful to understand and describe the interdependence of model parameters in the observation of practical non-identifiability, sequential optimization is entirely new.

Many fields are incorporating practical identifiability methods into modeling practices, primarily systems biology^55,56^ but increasingly in environmental models^57^, ecological population models^58^ and animal science^59^. However, the development of tools for practical identifiability analysis is still in its infancy and the number of reported data-based, interpretable cases is low^28^. As the role of digital twins in clinical decision-making, drug optimization, and disease diagnostics continues to grow^60^, reliable parameter estimation and uncertainty quantification will become increasingly important. Widespread application of practical identifiability still has hurdles, such as those of computational efficiency, benchmarking, and a priori prediction^28^. Our work aims to not only help address the issue of a priori prediction, but also serves as an exemplar of identifiability-driven practices in AD and other clinically-relevant computational models. Reliable model prediction and reporting of prediction uncertainty will be critical in closing the gap between model usage in the scientific domain and model usage in clinical decision-making. Modelers must be able to determine how biological and physiological assumptions can interrupt identifiability, and how to reconcile the importance of each facet of mechanistic modeling. Clinicians that use these models must be informed, whether by means of quantitative data attributes or qualitative patient phenotypes, under what circumstances we can expect the model to be dependable.

## Methods

### The ADBC Model

The ADBC model is described in detail in the literature^61,62^, and a mathematical proof of concept tested on real patient data can be found in our group’s previous work^15^. The ADBC hypothesis relies on well studied observations that the initiating event in AD is related to the formation of amyloid-*β* plaques in the brain caused by abnormal processing of the amyloid-*β* peptide. Accumulation of amyloid plaques can be observed through increased amyloid PET tracer retention. Abnormal phosphorylation, or hyperphosphorylation, of tau leads to its aggregation into paired helical filaments, which are the main components of neurofibrillary tangles, a biomarker for neuronal injury. After a delay, which varies across subjects, neural dysfunction and neurodegeneration dominate. Neurodegeneration can be observed through assessing brain volume or atrophy. In this work, only the biomarkers amyloid-*β*, phosphorylated tau, and neurodegeneration are considered. As shown in Equations 1-3, this cascade relationship is formalized through a system of ordinary differential equations:

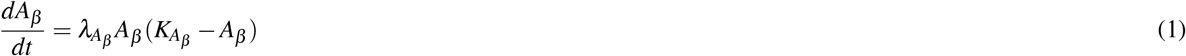

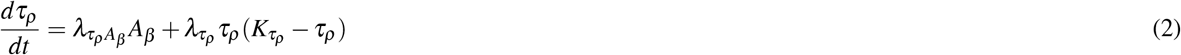

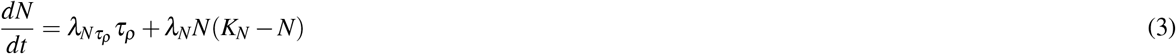

where 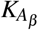 is the carrying capacity of amyloid, 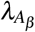 is the growth rate of amyloid, *A*_0_ represents initial amyloid-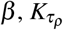 is the carrying capacity of tau, 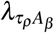 is the growth rate of tau driven by amyloid, 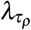 is the independent growth rate of tau, *K*_*N*_ is the carrying capacity of neurodegeneration, 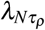 is the growth rate of neurodegeneration driven by tau, and *λ*_*N*_ is the independent growth rate of neurodegeneration. Only the initialization of amyloid-*β* is a parameter, while the initial values of phosphorylated tau and neurodegeneration are set to zero.

### Data sourcing and selection

Longitudinal PET data were obtained from the Alzheimer’s Disease Neuroimaging Initiative (ADNI) database (adni.loni.usc.edu), a large multinational study that has yielded key insights into the temporal dynamics of Alzheimer’s disease biomarkers^63,64^. ADNI began in 2004 and comprises five sequential studies—ADNI1, ADNIGO, ADNI2, ADNI3, and ADNI4—which followed subjects up to 15 years. To date these protocols have recruited over 1500 adults, ages 55 to 90, to participate in the research, consisting of cognitively normal older individuals, people with early or late MCI, and people with early AD. All amyloid and tau were obtained from PET scans and processed by Berkeley, where standardized uptake value ratios (SUVRs) were computed using a predefined region of interest or as an average across multiple regions. Only PET scans with a spatial resolution of 6 mm were included in the analysis. Amyloid-*β* burden was assessed using a summary SUVR defined as the weighted average of florbetapir uptake in the frontal, anterior/posterior cingulate, lateral parietal, and lateral temporal regions, normalized by the whole cerebellum. Amyloid scans were collected by the UC Berkeley lab from 1614 subjects spanning ADNI1, GO, 2, 3, and 4. Tau pathology was summarized using the weighted mean SUVR of the meta-temporal region from Jack et al.^65^, normalized to the inferior cerebellar grey matter. Tau scans were collected by Berkeley from 894 subjects spanning ADNI2, 3, and 4. Neurodegeneration was proxied by hippocampal volume normalized by intracranial volume (ICV), derived from MRI volumetrics collected approximately annually, using the scan closest in date to each subject’s PET exam. Hippocampal volume and ICV were obtained from 2172 subjects spanning ADNI1, GO, 2, and 3. To ensure adequate longitudinal coverage, only subjects with at least two exams per biomarker were included, resulting in a final cohort of 223 participants shared across these three datasets drawn from ADNI. Subjects were identified by their initial diagnosis in ADNI.

### Data preparation

As the disease progresses, amyloid-*β* and phosphorylated tau measurements increase, while hippocampal volume decreases. Each biomarker measurement was scaled such that the middle 90% of measurements fell into a fixed range between 0 and 1, which represent the theoretical minimum and maximum biomarker abnormality levels^15^. This was done by setting *X*_*max*_ equal to the 95th percentile threshold of values and *X*_*min*_ equal to the 5th percentile threshold such that

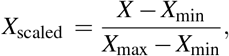

where *X* represents the original value and *X*_*scaled*_ represents the scaled value. The scaled values for hippocampal volume were subtracted from 1. In this way, all biomarker values would increase as the disease progresses.

### Parameter Estimation

For each equation, model parameters are optimized against subject-specific data from the ADNI by minimizing the objective function:

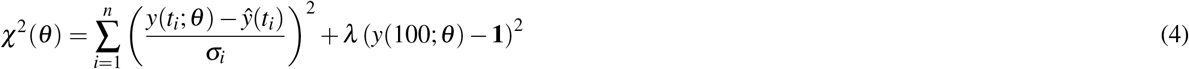

where *n* is the number of data points, *y*(*t*_*i*_; *θ*) denotes the biomarker trajectory values predicted by the model (Equations 1-3) at time *t*_*i*_, *ŷ*(*t*_*i*_) are the corresponding observed data points from ADNI, *σ*_*i*_ represent measurement errors, and *λ* is the constraint coefficient enforcing that the biomarker reaches its maximum at age 100.

The optimized parameter set *θ\** is found by numerically solving:

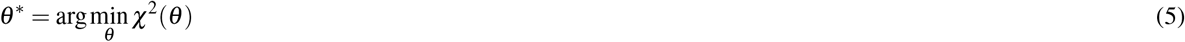

The relative measurement error for amyloid-*β* PET SUVR has been reported as approximately 3% of the median SUVR in cognitively normal (CN) controls, which is 1.38^66^. In our study, this error was estimated by calculating the median scaled amyloid value within the control group (on a 0 to 1 scale) and taking 3% of that value. We found that a 3% error was also reasonable for tau^67,68^ and neurodegeneration^69^, so this procedure was repeated for these downstream biomarkers. Consequently, the measurement errors *σ*_*i*_ used in the objective function (Equation 4) are set to 0.00414, 0.0182, and 0.0122 for amyloid-*β*, tau, and neurodegeneration, respectively.

Parameter estimation closely follows the approach in our previous work^15^, where optimization was performed iteratively with multiple initial guesses. Some key differences include the setting of the constraint coefficient *λ*, which is now tuned as a hyperparameter to improve model fitting. Additionally, all parameters, including those related to tau, were optimized using the Limited-memory Broyden–Fletcher–Goldfarb–Shanno (L-BFGS) algorithm^70^ within the bounds listed in Table 3. We assumed that amyloid-*β* aggregation is minimal at birth, and therefore imposed an upper bound of 0.2 on the initial amyloid-*β* value at age 0. Final personalized parameters correspond to the ODE system solution yielding the minimal loss. Fits were performed for all subjects.

### Identifiability analysis

First, structural identifiability analysis was completed with GenSSI, a software toolbox for structural identifiability analysis of biological models^71,72^. GenSSI uses a generating series approach and summarizes the non-zero elements of the Jacobian of the series coefficients in a tableau^73^. The output tableau for Equations 1–3 is in the supplementary, and shows that all parameters are globally identifiable.

Our practical identifiability analysis used the profile likelihood, a commonly used method for assessing practical identifiability^25,27–29,31,55,74^. While some authors recommend incorporating a range of tests to ensure confidence in results^43^, others note that the profile likelihood and its derivatives are the only methods that can effectively and reliability quantify parameter uncertainty and diagnose identifiability issues under discrete sampling procedures and measurement error^75^. Therefore, we resorted to applying only this technique to 65 of the 223 subjects. If measurement noise *σ*_*i*_ is from a Gaussian distribution, then

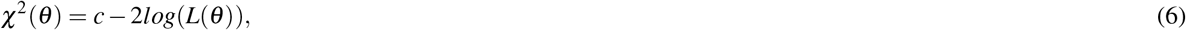

where *L*(*θ*) is the likelihood and *c* is a constant. Taking the objective function as a placeholder for the likelihood, we used the objective function to define the profile likelihood as

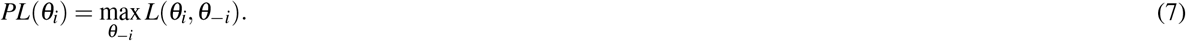

We then define likelihood-based confidence intervals by the confidence region

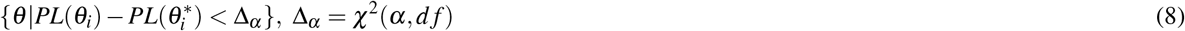

with *d f* = 1, since each profile likelihood is individually calculated for each parameter of interest^27^. Finally, we defined a practically identifiable parameter to be a parameter with a finite likelihood-based confidence interval, and a practically non-identifiable parameter to be one with an infinite interval. To perform a profile likelihood scan, we started at the optimal value of each parameter, and new parameter values were proposed in increasing and decreasing directions until the profile likelihood reaches the threshold 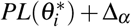. An infinite confidence interval is a consequence of only reaching the threshold on one end, or not at all. Two types of steps were used, logistic stepping for parameters (e.g. initial value for amyloid) of small magnitude, or linear stepping. Linear stepping followed a secant approximation for adaptive step size

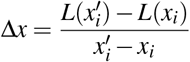

where 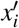 is the new proposed value for the parameter of interest, and *x*_*i*_ is the current value. A fixed step size is also given, and the minimum of the fixed step size and the adaptive step size was taken such that steps had consistently fine resolution. Scanning ended if the loss surpassed the given threshold for a subject or if an upper or lower bound was reached. Bounds for scanning were reduced to where the profile likelihood remains underneath the threshold, so the entire parameter space did not need to be searched.

### Uncertainty quantification

For uncertainty quantification, we chose to sample parameter values from their posterior distributions, as done in other studies^12^. Joint posterior distributions were found for parameters that were *identifiable*^30–32^ using the Metropolis-adjusted Langevin algorithm (MALA)^32,76^, a Markov chain Monte Carlo (MCMC) method for obtaining random samples from a probability distribution for which direct sampling is difficult. MALA uses a combination of two mechanisms to generate the states of a random walk that has the target probability distribution as an invariant measure: 1) the algorithm leverages Langevin dynamics, calculating the gradient of the target probability density function so that proposed values follow the gradient to where the distribution has higher density and 2) proposals are accepted or rejected using the Metropolis-Hastings algorithm, which guarantees convergence after some time. Multivariate normal distributions were used for the prior. Rather than sampling all parameters at once, we implemented a *coordinate* MALA that involved sampling through alternating parameters. For example, if both *K*_*N*_ and *λ*_*N*_ were identifiable and we sought to find their joint posterior distribution by iterating through a total of 5000 samples, we may first sample 500 values for *K*_*N*_, then 500 values for *λ*_*N*_, then 500 samples again for *K*_*N*_, etc. The number of samples prior to alternating was a hyperparameter we set for better convergence. Before sampling, all parameters were scaled to remain between 0 and 1. Given the bounds shown in Table 3, transformations were only necessary for *K* parameters and *A*_0_, while all other parameters (rate constants) were already between 0 and 1. We let *T* : Ψ→ [0, 1] be the transformation, where Ψ is the original parameters space. Then, for K parameters, where Ψ = [0, 10]:

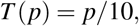

and for *A*_0_, where Ψ = [0, 0.2]:

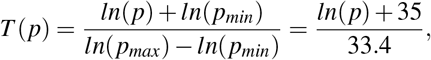

both of which used min-max scaling. Trace and autocorrelation plots were created to assess convergence. We took 1000 parameter samples from joint posterior distributions and simulated 1000 new solutions to Equations 1-3. Then, the mean and standard deviation of these trajectories were calculated to create graphs indicating where 95% of trajectories are expected to land. These averaged confidence intervals were calculated for 45 subjects (15 CN, 15 MCI, 15 AD).

The MALA method failed for the identifiable parameters of one biomarker equation in one subject (RID 6661), producing posterior distributions for neurodegenerative parameters that excluded the local minimum and proposed samples that increased without stabilization. This invalid posterior would then manifest as solutions that did not pass through any data points. To combat this, we implemented a new objective consisting of a local quadratic estimation of the initial objective function The gradient and Hessian of the initial objective function were found numerically using a first order central difference method, with discretization of 10^−11^. For this subject, the quadratic estimation was implemented as the new objective for the neurodegeneration equation in the calculation of the log likelihood for posterior sampling.

## Acknowledgements

JJ and JRP were supported in part by National Science Foundation (NSF) DMS2052676 and the National Institutes of Health (P30AG072958). WH was supported in part by NSF DMS2052685 and the National Institute of General Medical Sciences through grant 1R35GM146894. Data collection and sharing for this project was funded by the Alzheimer’s Disease Neuroimaging Initiative (ADNI) (National Institutes of Health Grant U01 AG024904) and DOD ADNI (Department of Defense award number W81XWH-12-2-0012). ADNI is funded by the National Institute on Aging, the National Institute of Biomedical Imaging and Bioengineering, and through generous contributions from the following: AbbVie, Alzheimer’s Association; Alzheimer’s Drug Discovery Foundation; Araclon Biotech; BioClinica, Inc.; Biogen; Bristol-Myers Squibb Company; CereSpir, Inc.; Cogstate; Eisai Inc.; Elan Pharmaceuticals, Inc.; Eli Lilly and Company; EuroImmun; F. Hoffmann-La Roche Ltd and its affiliated company Genentech, Inc.; Fujirebio; GE Healthcare; IXICO Ltd.;Janssen Alzheimer Immunotherapy Research Development, LLC.; Johnson Johnson Pharmaceutical Research Development LLC.; Lumosity; Lundbeck; Merck Co., Inc.; Meso Scale Diagnostics, LLC.; NeuroRx Research; Neurotrack Technologies; Novartis Pharmaceuticals Corporation; Pfizer Inc.; Piramal Imaging; Servier; Takeda Pharmaceutical Company; and Transition Therapeutics. The Canadian Institutes of Health Research is providing funds to support ADNI clinical sites in Canada. Private sector contributions are facilitated by the Foundation for the National Institutes of Health (https://www.fnih.org). The grantee organization is the Northern California Institute for Research and Education, and the study is coordinated by the Alzheimer’s Therapeutic Research Institute at the University of Southern California. ADNI data are disseminated by the Laboratory for Neuro Imaging at the University of Southern California.

## Author contributions statement

J.J. developed and implemented the code related to the identifiability methods, posterior sampling, data pre-processing pipeline, and post-processing analysis used in this study. J.J., J.R.P., and W.H were involved in the conception and design of the work, and the evaluation of the results. J.R.P. provided the clinical motivation, while W.H. provided the mathematical modeling expertise, and both authors supervised the work. J.J drafted the manuscript. All authors reviewed the manuscript.

## Ethics Declaration

### 0.1 Competing Interests

J.R.P. has served on medical advisory boards for cortechs.ai, Biogen and icometrix. No competing interest is declared for other authors.

## Notes

https://adni.loni.usc.edu/

