## Supplementary Material for "Identifiability-Guided Assessment of Digital Twins in Alzheimer’s Disease Clinical Research and Care"

August 17, 2025

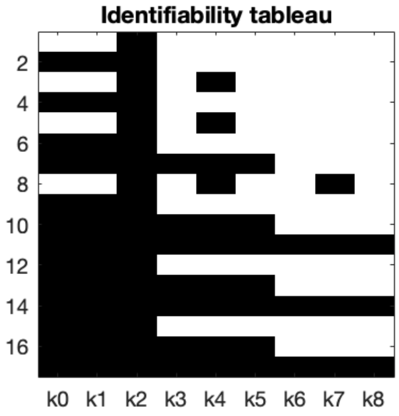


Figure 1: Identifiability tableau generated by GenSSI, with parameters "k0" through "k8" representing $K_{A_\beta}$, $\lambda_{A_\beta}$, $A_0$, $K_{\tau_\rho}$, $\lambda_{\tau_\rho A_\beta}$, $\lambda_{\tau_\rho}$, $K_N$, $\lambda_{N\tau_\rho}$, $\lambda_N$, respectively.

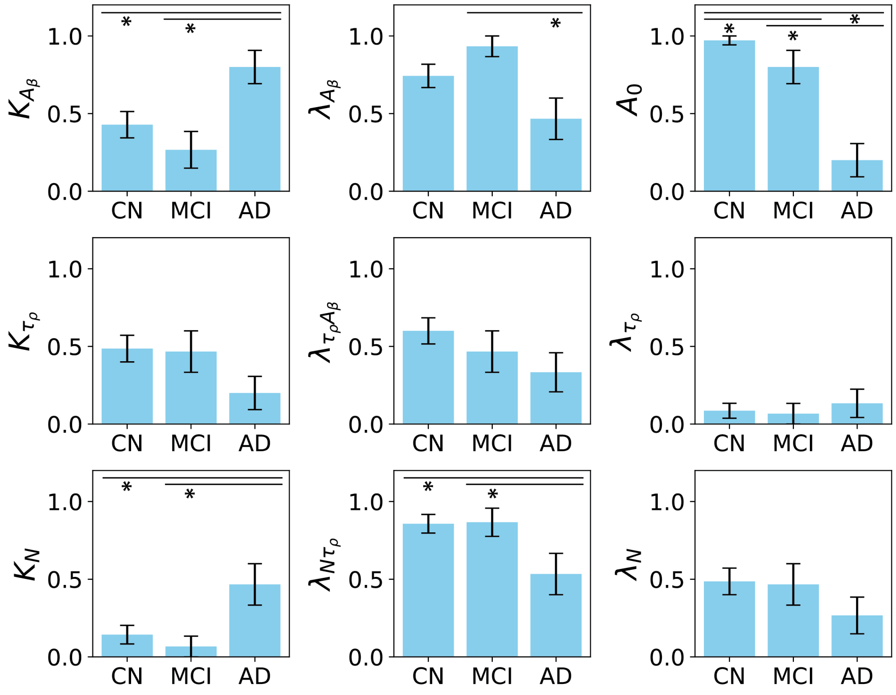


Figure 2: Bar graphs of average identifiability across disease groups. Horizontal bars indicate a post-hoc test, and astericks indicate significance.

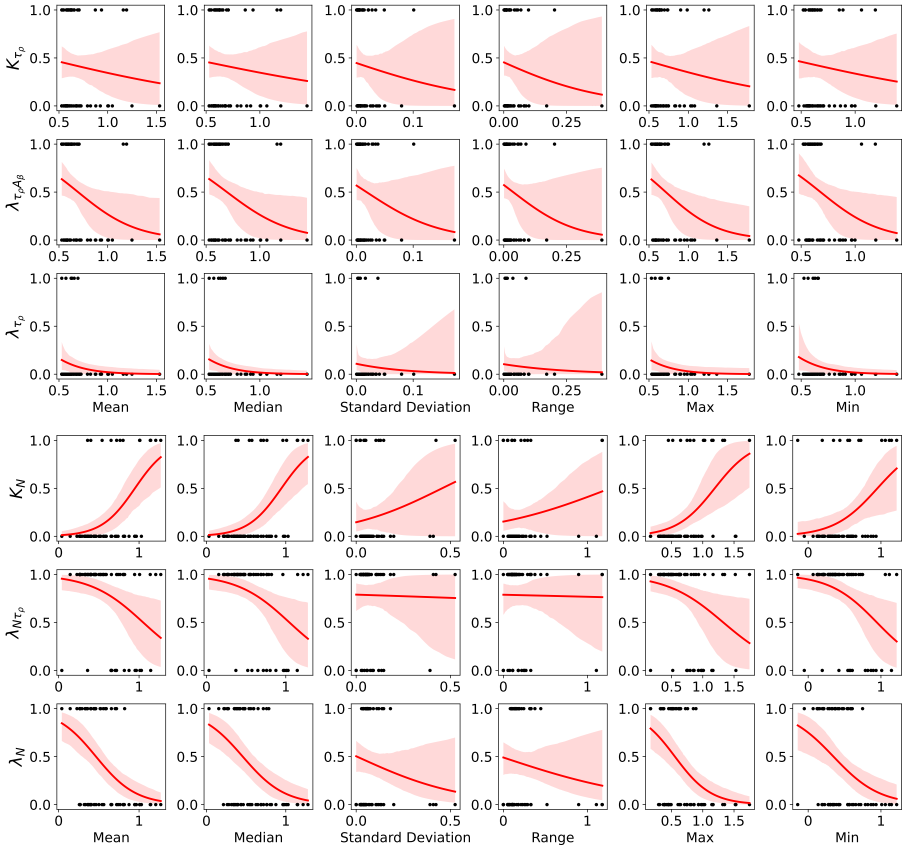


Figure 3: Logistic regression plots of the practical identifiability of the parameters in the phosphorylated tau and neurodegeneration equations against biomarker data metrics.

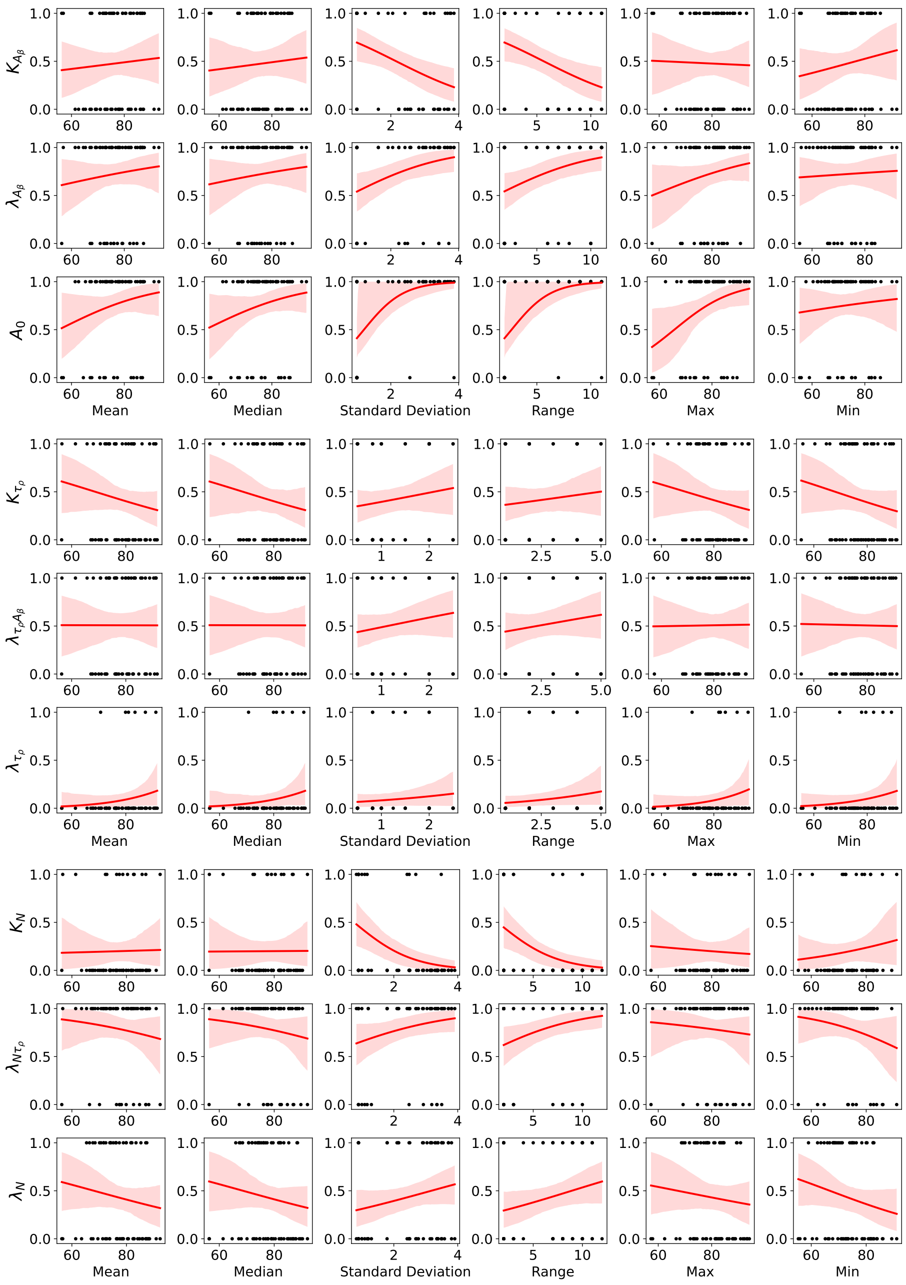


Figure 4: Logistic regression plots of the identifiability of all nine parameters in the amyloid, phosphorylated tau and neurodegeneration equations against age data metrics.

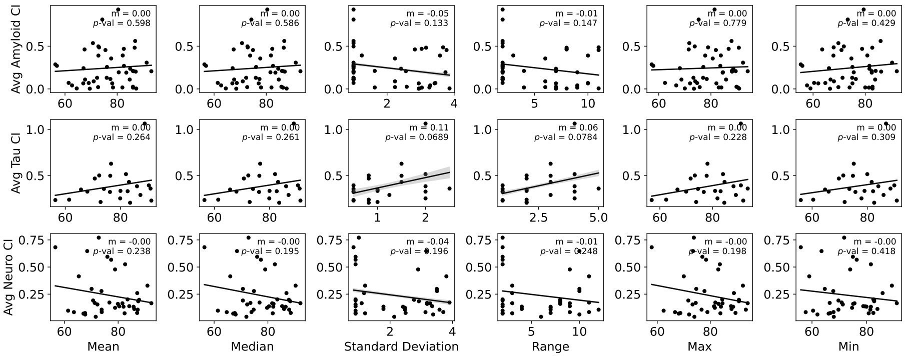


Figure 5: Confidence interval widths of predicted disease progression, averaged across examination dates, plotted against the mean, median, standard deviation, range, maximum, and minimum of the ages of examination.

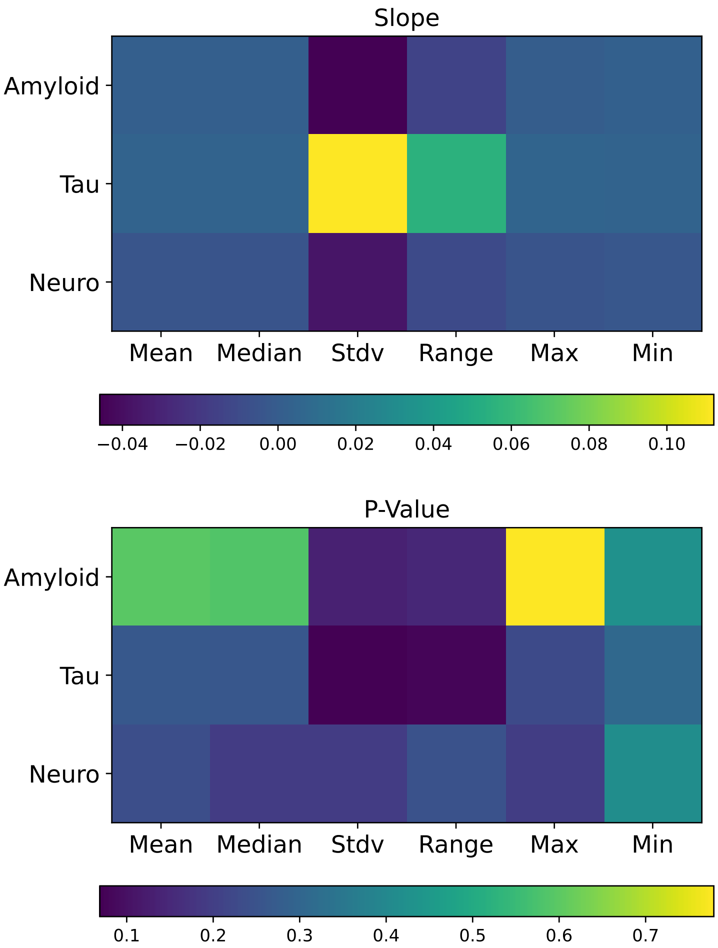


Figure 6: Slope and p-values obtained from a linear regression performed on each scatterplot in Supplementary Figure 5.
